# Non-segmented unsupervised learning of multispectral whole slide images for robust analysis of tissue repair and regeneration

**DOI:** 10.1101/2025.08.19.671136

**Authors:** Kody Paul Mansfield, Tamara Mestvirishvili, Bibi Subhan, Valeria Mezzano-Robinson, Dianny Almanzar, Sydney Hanson, Jimin Tan, Cynthia Loomis, David Fenyo, Aristotelis Tsirigos, Piul S. Rabbani

## Abstract

Analyzing whole tissue architecture remains challenging due to the inherent complexity of multicellular organization, variable morphology, and the limitations of conventional segmentation-based image analysis. Traditional approaches often rely on partial sampling or nuclear/cytoplasmic boundaries, which risk introducing bias and fail to capture the contextual interplay of diverse tissue compartments. To overcome these barriers, we developed a segmentation free framework for analyzing multispectral whole slide images (WSIs). By tiling WSIs into fixed sized regions and extracting quantitative tile features, we applied unsupervised machine learning to systematically reveal patterns of tissue organization at scale. This approach preserved spatial context without the need for cell-level delineation, recapitulating expected compartments such as epidermis, adipose, and scab, while also revealing subtle but coherent substructures within stromal and granulation regions. Applied to murine wound healing, the method distinguished wild type from diabetic repair dynamics without prior labels, uncovering both gross and nuanced differences in tissue composition. Together, this work establishes a robust, unbiased strategy for whole-tissue analysis that circumvents the limitations of segmentation, leverages unsupervised learning for discovery, and advances the study of tissue repair and regenerative pathology.

## INTRODUCTION

Tissue repair and regeneration is integral for the survival of complex multicellular organisms. Positioned at the interface between our body and the external environment, skin is frequently exposed to injurious stimuli and must therefore maintain robust regenerative capacity upon injury. To this end, the skin, our body’s largest organ, has evolved into one of the most regenerative tissues across the animal kingdom such as in *Drosophila*, zebrafish and chick embryos as well (Peña and Martin 2024; George and Martin 2022), and indeed pathologies of this process (such as the wounds often observed in diabetic skin) have poor health outcomes (Peña and Martin 2024; Armstrong et al. 2023). The skin’s remarkable repair and regenerative capabilities are underpinned by the skin’s complex tissue architecture and a multitude of spatiotemporally coordinated cellular and molecular mechanisms.

Skin is compartmentalized into three major tissue layers, from superficial inwards: the epidermis, dermis, hypodermis, and a fourth layer only in rodent skin, the muscle panniculus carnosus (**Figure 1**) (Zomer and Trentin 2018). The epidermis confers biomechanical and hydrophobic properties, forming a plastic cling wrap like water-tight seal (Zomer and Trentin 2018). Located beneath the epidermis, the dermis is a stromal tissue largely comprised of extracellular matrix and sparsely dispersed dermal fibroblasts, in addition to peripheral nerve endings and both blood and lymphatic vasculature (Mansfield and Naik 2020). The papillary dermal fibroblasts provide a supportive niche for the development and maintenance of the hair follicles while the reticular fibroblasts provide structural support by producing fibrillar extracellular matrix and during wound repair, express smooth muscle actin (Sennett and Rendl 2012; Philippeos et al. 2018). The hypodermis is an adipose tissue layer of lipid-filled adipocytes (Treuting et al. 2012). Unlike humans, lower-level mammalian species, such as mice, have a pronounced layer of striated muscle as the deepest layer of skin. This tissue, while typically responsible for skin twitching and movement, plays a key role in rodent wound healing models as the muscle contracts to significantly reduce wound size and enacts different molecular mechanisms than human skin wound closure. Each of the of tissue layers comprising skin have to coordinate to achieve repair and regeneration.

**Figure 1.**
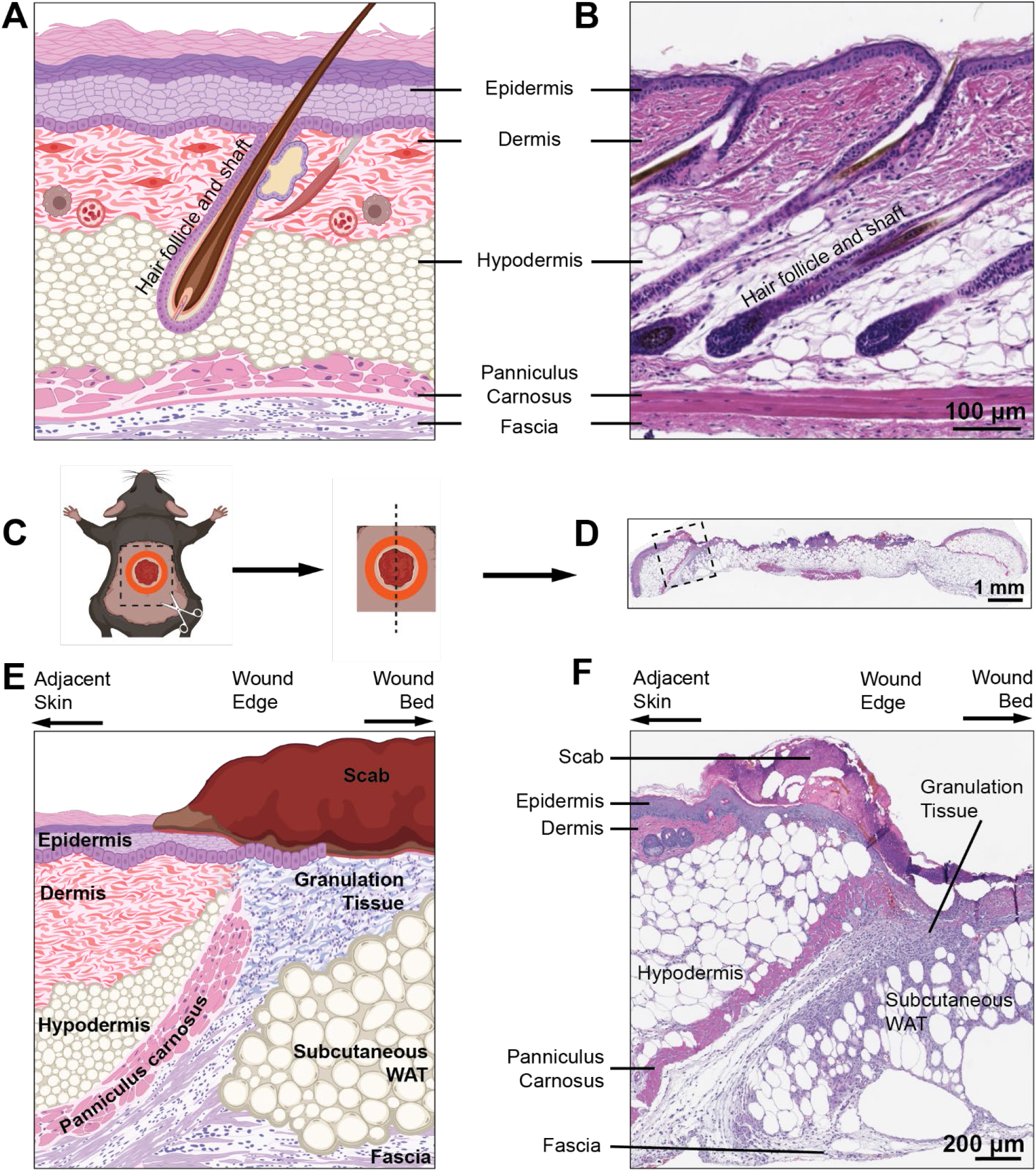
Mouse skin architecture and wound sample orientation. A) Schematic of intact mouse dorsal skin in B. C) Dorsal full-thickness (all tissues above fascia excised) wounded skin bisected along cranial-caudal axis to generate sections spanning the wound (D). This section is H&E-stained. E) Schematic of excisional/full-thickness wound section microscopic image in F. This is one end of the wound, as shown in the dashed box in D. F) Wound section from a LepR^db/db^ mouse dorsal wound.

Following a wound in the skin, a new transient structure termed granulation tissue begins to form at the wound edges through an intensely coordinated process among fibroblasts, blood vessels and infiltrating immune cells (Figure 1) (Peña and Martin 2024). Together these three cell populations systematically aid one another to progress the granulation tissue deeper into the wound bed. Epidermal cells located adjacent to the wound, migrate into the wound on the scaffold provided by the granulation tissue to close the tissue and seal off the wound bed, known as clinical re-epithelialization. Wound healing continues past re-epithelialization in attempt to reinstate the functional barrier roles of skin. The highly spatiotemporally regulated wound healing events are disconnected in non-healing and chronic wounds, such as in diabetes, venous disease, peripheral arterial disease.

Imaging is the convergence point for the different disciplines that study skin repair and regeneration. Advanced microscopy of multi-color immunostaining, have informed our understanding of the multi-tissue collaboration in skin and divergence in disease/disorder or genetic preclinical wound healing models and clinical cases. Building on this foundation will require systematic, high-resolution imaging of both typical acute healing and chronic, non-healing wounds. Such studies are paramount to define the spatiotemporal dynamics of normal repair, pinpoint where these programs break down in diabetes, and ultimately inform clinical treatments.

Prior to the development of digital whole-slide scanning, the only way to survey an entire microscope slide was through visual inspection at the microscope. High magnification is required to obtain high resolution images at a cellular level within each tissue, but this inherently limits the field of view that is observed and that can be captured within a single image. As a result, traditional microscopy images capture only a small fraction of a given tissue within a large biological specimen. This constraint impedes applicability of traditional microcopy, to manual image acquisition as it is impractical to build complete image datasets spanning even a single tissue type present in an organ, much less the entire specimen across a whole slide. Instead, imaging-based studies are limited to the specific region of tissue which the researcher manually chooses to capture. The utility of microscopy as an experimental approach can be diminished, as errors can be inadvertently introduced through sampling bias during this traditional image acquisition process.

Manual microscope photography has significant limitations when it comes to capturing comprehensive data about regions where multiple cell types interact, such as stromal tissues in the skin, dermis, and granulation tissues. Traditional analyses often rely on binary combinations of markers with counterstains to build data on cell-cell relationships, which can result in a loss of context regarding the tissue’s environment. This approach does not facilitate the discovery of new patterns of expression or regions of difference. Although recent advances in large-scale multi-marker immunostaining of protein or RNA probes have opened up new avenues for exploration, they remain constrained to individual photographs, such as those produced with tools like Cell Pose. A critical limitation in assessing complex organs like skin is the insufficiency of segmentation techniques to identify and collect data on all cell types involved in tissue repair and regeneration. Figure 2 demonstrates a region of interest in the granulation tissue of a murine wound sample, immunostained for F4/80, a pan macrophage marker, and CD31, an endothelial cell marker, counsterstained with DAPI to identify nuclei. Neither cell type immunostained has textbook schematic-type round cell shapes and in fact may present in a section without a nuclei present, such as the macrophages highlighted in Figure 2A. Whether in the wound adjacent intact dermis (Figure 2B-C) or wound granulation tissue (Figure 2B,D), a segmentation approach would not account for these cell types due to the lack of nuclei in the imaging plane.

**Figure 2.**
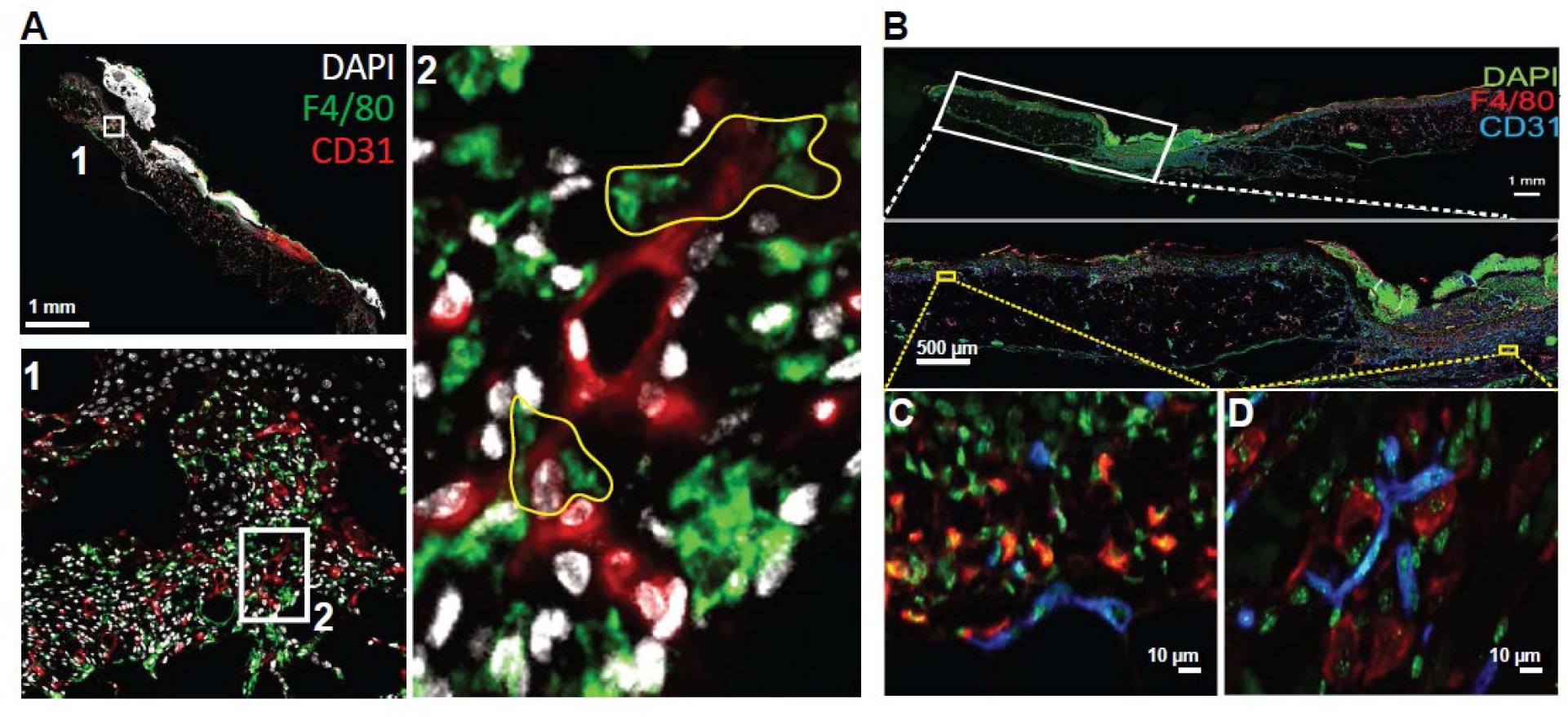
Limitations of segmentation in image analysis. A) Two antibody immunostained and counterstained fluorescent image demonstrating green fluorescent F4/80+ cells (yellow outline) that do not have a visible nucleus in this wound section. The nuclei in proximity belong to other cells, not the outlined F4/80+ macrophages. B) Second example of limitations of nuclear segmentation in identifying cells, comparing two stromal tissue, wound adjacent dermis (C) and the granulation tissue (D).

To address these challenges in our methodology, we have developed a segmentation-free approach for analyzing WSIs. This methodology eliminates the need for delineating nuclear or cytoplasmic boundaries, enabling direct quantification of tissue features using intensity-based and spatial metrics. Such an approach is particularly advantageous for tissues exhibiting irregular morphologies or partial cell visibility. Our strategy preserves the anatomical context across gigapixel-scale images, facilitating comprehensive quantification of entire tissue sections. Moreover, our workflow is fully compatible with the large file formats generated by whole-slide scanners, which often exceed several gigabytes in size. These formats typically employ a pyramid structure, allowing efficient navigation and analysis across multiple resolutions. By adopting this segmentation-free methodology, we circumvent the complexities and potential inaccuracies associated with traditional segmentation processes. Subsequently, we apply machine learning algorithms to enhance our understanding of the tissue sections, enabling the extraction of complex patterns and relationships within the data. Utilizing traditional machine learning (ML) algorithms offers several advantages, particularly when analyzing complete datasets. By focusing on a limited number of relevant features, these models reduce the risk of overfitting, as they are less likely to capture noise or irrelevant patterns. This simplicity enhances model generalization and interpretability. Moreover, traditional ML methods require less computational power compared to deep learning approaches, making them suitable for environments with limited resources. Additionally, traditional ML can be effectively implemented even without extensive training data, leveraging domain knowledge to adapt the model to the dataset. This approach eliminates the need to interpret missing data, as the focus is on complete, high-quality datasets, streamlining the analysis process. Consequently, traditional ML provides a practical, efficient, and interpretable framework for analyzing complete datasets without the complexities associated with deep learning.

## METHODS

### Murine Wound Model

All animal protocols were approved by the New York University School of Medicine Institutional Animal Care and Use Committee. C57BL/6N wildtype mice (Model Number: B6-M/F) were obtained from Taconic Biosciences (Germantown, NY, USA). Type 2 diabetic, *Lepr*^*db/db*^ mice (Strain Number: 000642) mice were obtained from Jackson Laboratory (Bar Harbor, ME, USA). The mice were anesthetized using 2% isoflurane (Henry Schein, Inc., Melville, NY, USA. Catalog number: 1182097), and the footpad pinch test was used to confirm that the mice were completely sedated. We used a well-established excisional mouse wound model as in our prior work (Subhan et al. 2021; Rabbani et al. 2018; Subhan et al. 2024). The hair was shaved from the mouse dorsum and Nair hair removal cream (Amazon. Model Number: 7784911) was used to remove remaining hair shafts. 10-mm-diameter full-thickness wounds were created using Acu-Punch Biopsy Punch (Fisher Scientific. Catalog Number: NC9236770). To prevent the panniculus carnosus from contracting the wound and resulting in premature closure, the wound was splinted with a 0.6-mm-thick silicone stent with an inner diameter of 10 mm and an outer diameter of 20 mm (using silicone sheets from Grace Bio-Laboratories, Bend, OR, USA. Catalog Number: 665581). Silk 4–0 braided reverse cutting suture (Henry Schein, Inc., Melville, NY, USA. Catalog Number: 1006830) was used to secure the stent to the skin surrounding the wound. To minimize scratching, chewing, and biting of the sutures by the mice, an occlusive adhesive dressing (Henry Schein, Inc., Melville, NY, USA. Catalog number: 777-9152) with a 12 mm window overlying the silicone stent was applied. The window allows air to exchange in the wound, while covering the actual sutures. Buprenorphine (Covetrus, Elizabethtown, PA. USA. Product Number: 059122) was administered for analgesia for 3 days post-operatively.

### Sample Processing

We dissected wounds from the mouse dorsum and then bisected along the rostral/caudal axis, such that the bisected edges are perpendicular to the block sectioning plane. This method ensures that we analyze samples from the center of the excisional wound. Immediately following dissection, we immersed skin wound samples in 4% paraformaldehyde (Electron Microscopy Sciences, Hatfield, PA. USA. Catalog Number: 15714) and crosslinked for 48 hours at 4ºC to preserve tissue structure and prevent degradation. The fixed wound samples underwent dehydration through a series of ethanol solutions of increasing concentrations starting at 70% ethanol for 60 minutes, 90% ethanol for 45 minutes, then two changes of 100% absolute ethanol, each for 45 minutes. We then used xylene immersion, 3 cycles of 30 minutes each as a clearing agent to replace the ethanol and further prepare the tissue for paraffin infiltration. We then embedded the wounds using molten paraffin wax at 56ºC, such that the wax solidifies at room temperature (RT) and provides structural support for sectioning. The paraffin-embedded blocks are cut into 5 μm sections using a microtome and mounted onto positively charged glass slides for further processing. Sample details are included in Extended Data Table 1.

### Histological Staining and Imaging

Each slide was stained with hematoxylin and eosin. These slides were imaged using Leica AT2 brightfield whole-slide scanner at 20X which produced whole slide image scans output as Aperio SVS files (.svs) (Figure 3A). This allowed for validation of sample structural integrity and initial histological analysis before proceeding to immunofluorescence staining.

**Figure 3.**
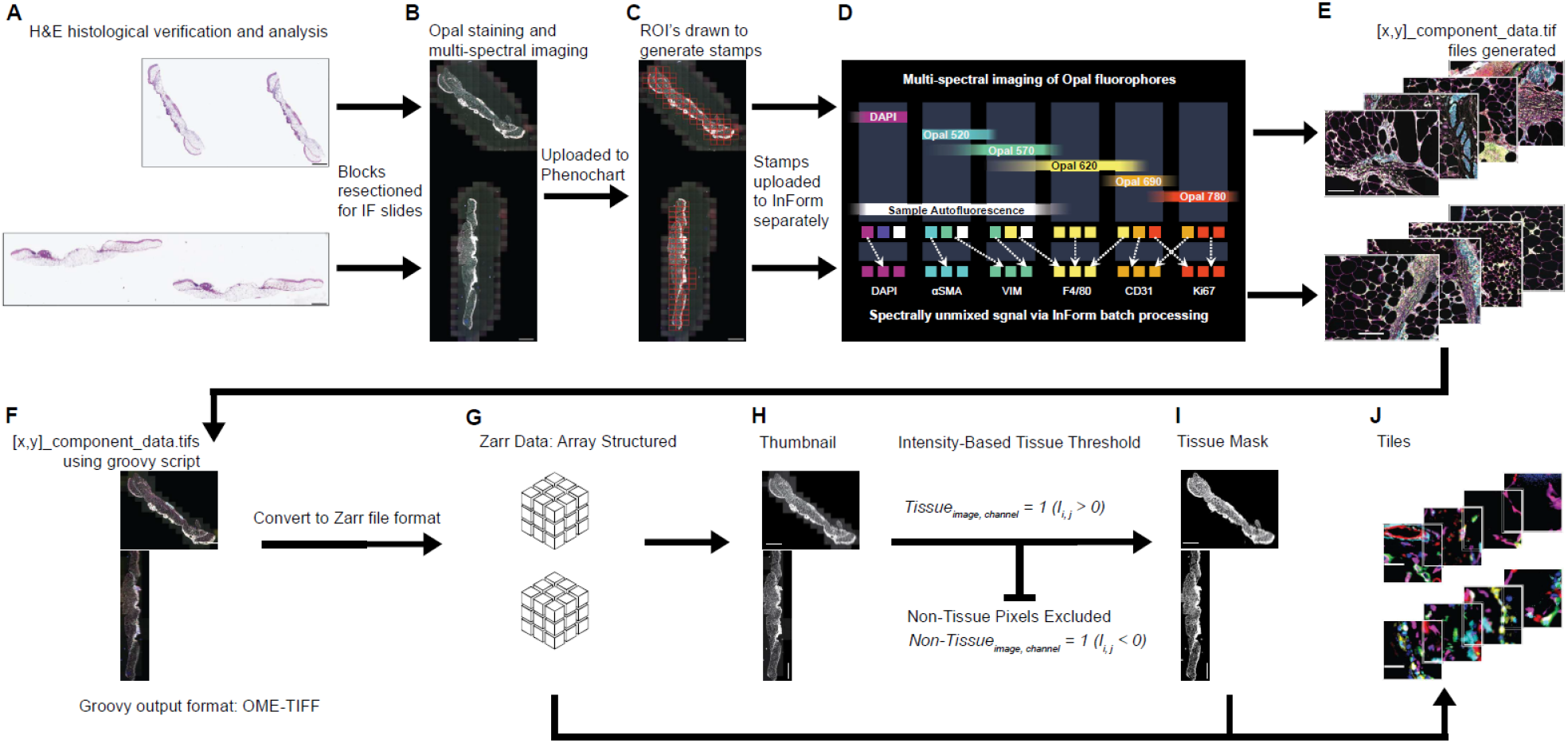
Whole slide imaging file preparation. (A) H&E-stained wound sections used for histological verification prior to multiplex imaging. (B) Opal multiplex immunofluorescence staining and multispectral image acquisition. (C) Regions of interest (ROIs) drawn in Phenochart to generate stamp images for each wound section. (D) Stamp images imported into InForm for algorithm-based batch processing and spectral unmixing to separate individual fluorophore channels. (E) Generation of [x,y]_component_data.tif files containing spectrally unmixed signals for each channel. (F) Stitching of unmixed component TIFFs into a pyramidal OME-TIFF using QuPath with positional metadata from TIFF tags. (G) Conversion of OME-TIFF to Zarr array format (CxHxW) with corresponding channel metadata. (H) Low-resolution thumbnail generation from Zarr slide. (I) Creation of binary tissue masks using an intensity-based threshold to exclude non-tissue pixels. (J) Division of masked tissue regions into fixed-size tiles with recorded coordinates for downstream analysis. Scale bars: (A-C, F, H-I) 2 mm; (E) 200μm; (J) 20μm

Then we sectioned wound sample paraffin blocks again and mounted 2 distinct samples per slide (Figure 3; Extended Data Table 1). Slides underwent the following processing: baking at 65°C for 1 hour, de-paraffinization with xylene (3 × 10 minutes), and rehydration through a graded series of ethanol solutions, 100% 1 × 10 minutes; 95% 1 × 10 minutes; and 70% for 10 minutes, followed by rinsing in 1x PBS and further fixation in 10% neutral buffered formalin for 20 minutes. All further processing for immunostaining occurred in a humidified chamber. We then incubated slides in antigen retrieval buffer (Akoya Biosciences Product Number: AR600250ML) and microwaved for 45 seconds at 100% power then for 15 minutes at 20% power, and finally rinsed with distilled water followed by TBST. Then the wound sections underwent incubation in blocking buffer (PerkinElmer, Catalog Number: ARD1001EA) for 10 minutes at RT. Following removal of the blocking buffer, we incubated the slides with the first primary antibody and incubated overnight 4°C temperature. After 3 × 2 minute washes in TBST at RT, we incubated slides with secondary antibodies which provide local horse radish peroxidase (HRP) activity using Polymer HRP Ms + Rb (Akoya Biosciences Product Number: ARH1001EA) for 10 minutes at RT. After washing, we added the Opal Fluorophore Working Solution at room temperature for 10 minutes to add fluorescent signal. To strip the primary-secondary-HRP complex, we microwaved slides for 45 seconds at 100% and 15 minutes at 20% power in antigen retrieval buffer. Once the slides cooled, we washed in TBST and repeated blocking, primary antibody binding, secondary antibody binding, fluorescent signal deposition, and antibody stripping for each primary antibody. Sequential staining order, target antigen markers, primary antibodies, and associated fluorophores are detailed in Extended Data Table 2. The power of this sequential staining, deposition of fluorescent signals, and stripping protocol enables multiplexed targeting of antigens using primary antibodies raised in the same host. Finally, we counterstained slides with Spectra DAPI solution (Akoya Biosciences Product Number: FP1490A) for 5 minutes at RT and mounted with coverslips using mounting medium. To image the whole slides, we used the Akoya Biosciences Phenoimager HT multi-spectral imaging system (previously named Vectra Polaris) at 20X which produced whole slide image scans output as PerkinElmer Vectra QPTIFF files (.qptiff) (Figure 3B).

### Spectral Unmixing

This imaging system uses multispectral imaging, enabling more precise signal detection compared to traditional microscopy, which relies on filters to separate fluorescent signals. Such filters limit the range of signals that can be captured in an image. As a result, the raw QPTIFF files require further processing to unmix the fluorescent signals.

QPTIFFs were opened in Phenochart, a digital pathology visualization and analysis tool developed by Akoya Biosciences. Using this software, we drew a region of interest (ROI) around each wound section to create a grid of image stamps. These stamps were then imported into *inForm*, Akoya’s image analysis software for multiplexed tissue imaging. We repeated this process for each wound section on the slide to obtain a single image for each wound sample. *InForm* applied algorithm-based batch processing to the stamps, generating a series of component_data.tif files for each wound sample with spectrally unmixed fluorescent signals (Figure 3C-E; Extended Data Files 1 & 2).

### TIFF to Pyramid

To reconstruct a single high-resolution image for each wound sample, the component_data.tif files, each representing a single field of view, were stitched into a pyramidal OME-TIFF using *QuPath* (v0.5.x) and the publicly available script *Convert TIFF fields of view to a pyramidal OME-TIFF* (Bankhead 2024). The script reads positional metadata from the baseline TIFF tags to accurately place each field within the stitched image. We used lossless compression to preserve fluorescence intensity values. The workflow involved selecting all unmixed component TIFFs for a sample, extracting positional metadata (X/Y coordinates, resolution, and size), building a sparse image server in *QuPath*, and then pyramidalizing the composite image into a multi-resolution OME-TIFF for efficient viewing and downstream analysis (**Fig 3F**).

### TIFF to Zarr to Tiles

We adapted the image preprocessing pipeline described by (Tan et al. 2025) which was originally developed to generate fixed-size tiles from IMC data for model training, to operate on whole-slide multiplex images. Multiplex TIFF files were converted to Zarr (C×H×W) with channel names recorded in a companion channels.csv (**Fig 3G**). From the Zarr slide, we generated a low-resolution thumbnail by block-reducing across channels and space using a scale set by the tile size (scaling_factor = tile_size/4) (**Fig 3H**). Intensities above the 95th percentile were clipped and values normalized to [0,1]. A binary content mask was obtained by thresholding the thumbnail (τ = 0.2) (**Fig 3I**). The mask was then resized with nearest-neighbor interpolation to a grid defined by a fixed pixel tile size (e.g., 128×128 px). Given the measured pixel size of 0.496 µm/px in both X and Y, each 128-px tile corresponds to ∼63.5×63.5 µm, closely matching the 64-µm tiles described in the original pipeline. Tiles were defined as grid cells with mask = 1; for each, we recorded the top-left pixel coordinates (h*tile_size, w*tile_size) and saved them to a positions file (**Fig 3J**) We also exported PNGs of the thumbnail, binary mask, and tile grid for QC. This procedure removes partial tiles at the image borders.

### Analysis

Features are extracted from whole-slide image data by focusing on the density of positive non-zero values within each tile. The pixel density for each channel (excluding the autofluorescence channel) is calculated by summing the non-zero-pixel values within each defined tile region. Tiles with non-zero densities are considered, while blank tiles are ignored. This generates a feature array, where each entry corresponds to the density of measurement for a tile in each channel. The approach leverages a density threshold to ensure that only meaningful tile regions contribute to the feature set. This method efficiently captures spatial patterns across multiple channels, providing insights into the structure and content of the slide image. In addition, we calculated the mean intensity of the channels per tile of each sample. We used a similar methodology to capture the positive tile density. We calculate the mean of each channel by utilizing the area of the tile and the wavelength intensity post normalization of each channel as features.

We created an interactive visualization that overlays spatial tile data onto a resized microscopy image, enabling precise alignment and inspection of computational results in their spatial context. It begins by loading tile metadata from a csv file and filters it to match a specified sample name. The corresponding whole slide image is then loaded and resized according to a user-defined scale factor. To align the tile coordinates with the image, the function calculates scaling factors based on the original and resized image dimensions and applies optional horizontal and vertical shifts to adjust the coordinate origin. Further adjustments to the scatter plot alignment can be made through additional scaling parameters in the x and y directions. The tile coordinates are transformed accordingly and plotted as semi-transparent black points using *Plotly* (v6.2.0x), with hover labels indicating each tile’s index. The resized image is set as the background of the interactive plot, and the result is saved as an HTML file for exploration. This method provides a visual tool to assess spatial patterns and validate tile-level data within the context of the original image, supporting downstream analysis and interpretation.

## RESULTS

### Tiling-based segmentation of multispectral images

Wound healing phenotypes between wild type (normoglycemic) and LepR^db/db^ (hyperglycemic, type 2 diabetes) mice, particularly on excisional or full-thickness skin wounds are significantly different, (p < 0.0001, unpaired t-test) (**Fig 4A**). Wound healing in the LepR^db/db^ model is severely impaired, requiring more than a two-fold increase in healing time of that of WT mouse wounds. This is a well-established preclinical wound model and the disparities in wound closure times between the two genotypes are also well documented (Galiano et al. 2004; Subhan et al. 2021; 2024; Rabbani et al. 2018). The delay in LepR^db/db^ mice is evident in intermediate time points in histology of wound sample sections (Subhan et al. 2024) and the conspicuous extreme differences are ideal for developing this image analysis tool to ensure applicability with a wide range of biological phenotypes.

**Figure 4.**
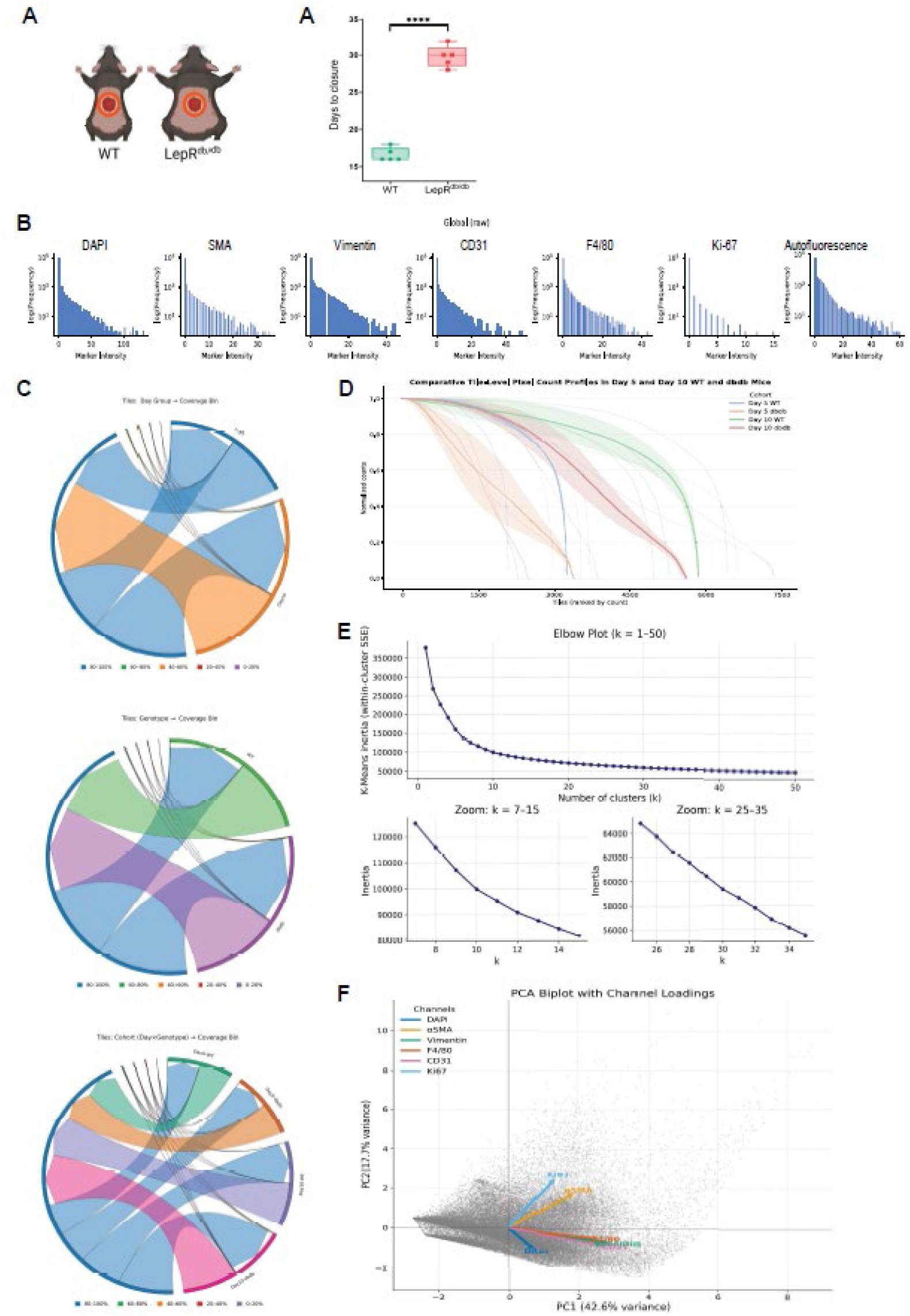
Global raw intensity and tile coverage across cohorts. (A) Log-scaled histograms of raw per-pixel fluorescence intensity for each channel (DAPI, αSMA, Vimentin, CD31, F4/80, Ki67, autofluorescence) across all cohorts. (B) Chord diagram showing the distribution of tiles by day group (Day 5 vs Day 10) across tissue-coverage bins (0–20%, 20–40%, 40–60%, 60–80%, 80–100%). (C) Chord diagram showing the distribution of tiles by genotype (WT vs db/db) across tissue-coverage bins. (D) Chord diagram showing the distribution of tiles by cohort (day × genotype) across tissue-coverage bins. E) Ranked normalized tile-level pixel counts by cohort, showing pre-clustering differences in spatial coverage.

Across all cohorts, raw fluorescence intensity distributions for each marker exhibited a steep peak near zero, with the shape and spread of the high-intensity tail varying by marker (**Fig. 4B**). Nuclear (DAPI), CD31, and autofluorescence channels displayed broader tails, indicating either strongly labeled tissue structures or greater staining variability. SMA, F4/80, and Ki67 had shorter dynamic ranges, with most pixels concentrated at low or moderate intensities, consistent with lower signal abundance or dimmer fluorophores, while Vimentin showed an intermediate profile with a gradual decline in mid-range frequencies. The autofluorescence channel, in particular, exhibited a long, diffuse tail without discrete high-intensity modes, reflecting non-specific background signal distributed throughout the tissue. Differences in maximum intensity across channels likely reflect both biological factors (marker abundance, tissue distribution) and technical factors (fluorophore brightness, spectral overlap, staining uniformity).

Chord diagrams of tile coverage by day, genotype, and combined cohorts (**Fig. 4C**) showed that most analyzed tiles across all groups fell within the 80–100 % tissue-coverage bin when coverage was defined from the union of non-zero pixels across all channels. This confirms that the tiling and masking workflow of our whole slide images consistently yields tiles that are largely filled with tissue signal, providing a robust foundation for subsequent density and mean intensity-based feature extraction. While this binning demonstrates overall uniformity in overall tissue presence, it does not capture variation within the top coverage category; these subtle differences, such as how close tiles are to actual complete coverage or how coverage is actually distributed across different channels, are resolved in the ranked curves (**Fig. 4D**).

Ranked normalized tile-level pixel count curves (**Fig. 4D**) revealed pre-clustering differences in spatial coverage across cohorts. For each sample, tiles were first ranked from highest to lowest tissue coverage, where coverage was calculated as the number of non-zero pixels (across all channels except autofluorescence) in the tile. The Y-axis values were then normalized by dividing each tile’s count by the maximum count observed in that sample, so that the most tissue-rich tile in each sample is scaled to 1.0 and all others fall between 0 and 1. This normalization allows curves from samples with different absolute tile counts or staining intensities to be directly compared on the same scale.

Curves with a slower decay, exhibiting a shallower or more gradual slope, such as those from Day 10 db/db, indicate that many tiles retain coverage values close to the sample maximum well beyond the top ranks. This pattern suggests a greater proportion of tissue-rich tiles, potentially linked to larger wound areas, thicker tissue layers/adipose tissue, or slower resolution of the wound. In contrast, faster-decaying curves with a steeper slope, Day 5 WT, drop sharply after the top-ranked tiles, reflecting more localized tissue coverage (a small group of tiles contributing disproportionately to overall coverage) and possibly indicating more advanced or complex wound closure with smaller residual tissue regions. Within the WT cohort, the Day 5 samples show a steeper slope than Day 10, suggesting that in this dataset, WT wounds have notable wound healing (in progress) phenotypes at POD5 samples, whereas by day 10 the samples tissue contains more healed and intact wound adjacent areas. These systematic differences in tissue abundance and spatial distribution are already apparent before clustering and provide an underlying spatial context that may influence how cohorts separate in feature space during downstream analysis.

After looking at the principal component analysis (PCA) on per –tile marker densities (non-zero pixel counts of all channels excluding the autofluorescence) after z-scaling. The PC1-PC2 plot shows each tile as a gray point, arrows indicate the loading vectors for each channel (**Fig. 4F**). The length of the arrow represents the contribution of the channel to the plotted variance, and the angle between the arrows indicates the correlation. The tiles form a continuous manifold rather than obvious sections, indicating smooth gradients in tissue composition across the tiles, consistent with the lack of a single, obvious elbow in the k-means WCSS curve (**Fig. 4E**). Overall, the PCA shows that the between-tile/sample heterogeny within the phenotypes is substantial and possibly organized along biologically interpretable axes rather than sharply separate clusters.

### Tissue Annotation with biological compartments

Taken together, the distributions, coverage profiles, and PCA manifold from Fig. 4 indicate substantial but continuous variation rather than sharply separated groups. Accordingly, and consistent with the absence of a clear elbow in the k-means curve, we did not use the elbow plot to fix the number of clusters. Instead, we tested whether the clusters determined by the unsupervised clustering aligned with known anatomical tissue compartments in the wound. We aimed to verify the classification/assignment of tile clusters against their anatomical tissue location in skin wound. We annotated the scanned images for the anatomically distinct skin wound tissue categories that constitute a skin wound – dermis, epidermis, granulation tissue, hypodermis, panniculus carnosus muscle, subcutaneous fat pad, fascia and scab (Treuting et al. 2012). We generated an interactive visualization of the autofluorescence (AF) image for each scanned section, with tile coordinates overlaid and adjusted to align with the underlying image (Fig. 5A). Using a lasso tool, we drew bounding regions to denote and color-code each tissue category (Fig. 5B–D). The resulting annotations were exported to a CSV file for downstream use. To verify accuracy, we overlaid the color-coded tissue boundaries onto the multispectral image of the same section (Fig. 5E–G)

**Figure 5.**
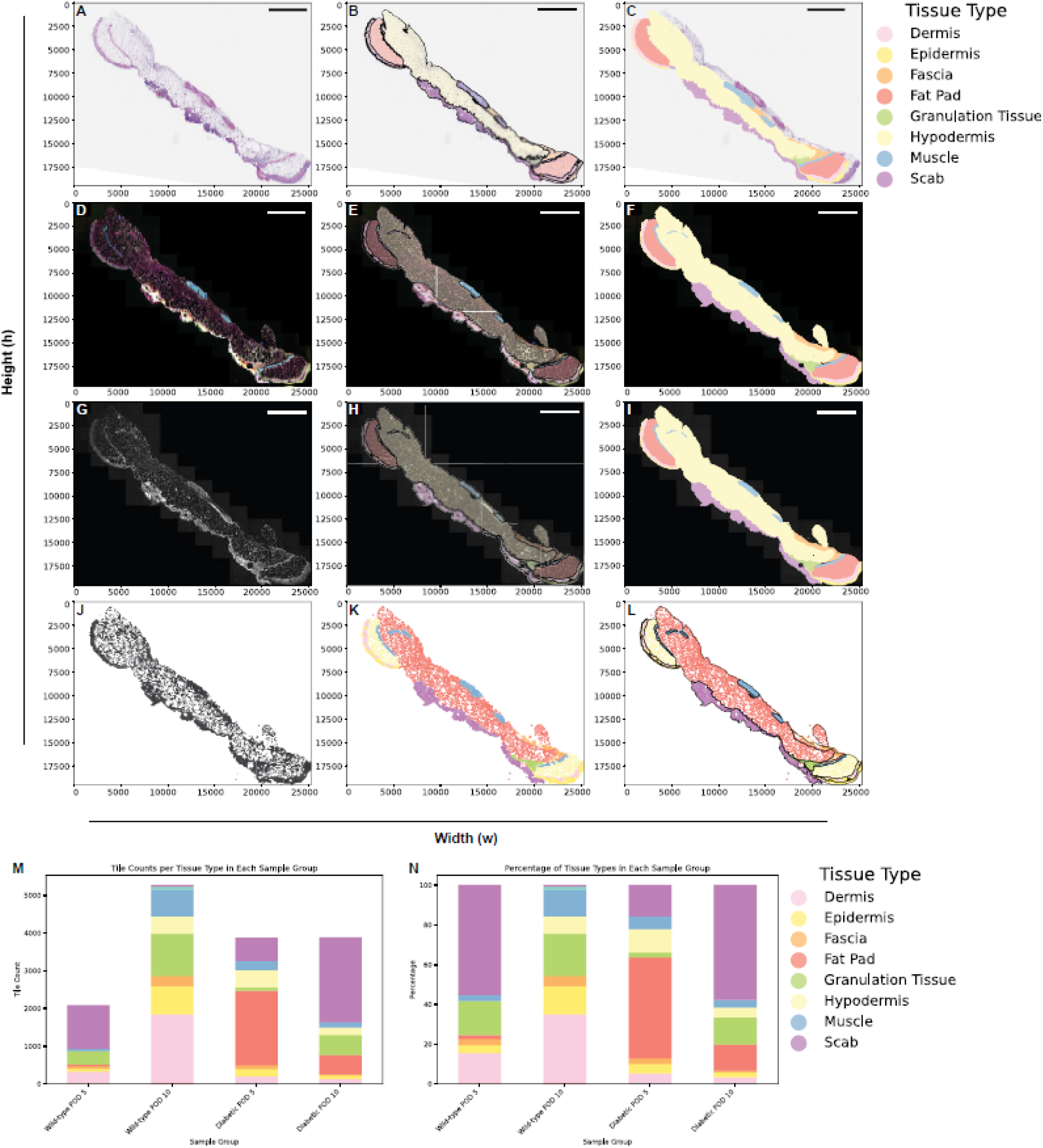
Assignment of tiles to user-defined anatomical tissue type categories. (A) Workflow of annotating tiles by skin wound tissue types. (B) Scanned H&E image of a wound section. (C-D) User-defined annotation of boundaries of broad anatomical tissue categories using lasso tool, overlaid on H&E image, and color-filled by category. (E) Multispectral image of wound section in (B). (F-G) Annotated color-coded tissue categories overlaid on (E). (H-J) Opal autofluorescence channel only image of with overlaid tissue categories. (K) Tile position map. (L-M) Tile map from (K) colored by the user-defined tissue type category annotations. (O) Tile counts per tissue category across the 4 biological conditions in this study. (P) Percentage of tissue categories in each sample group, the 4 biological conditions.

We annotated the scanned images for the anatomically distinct skin wound tissue categories that constitute a skin wound – dermis, epidermis, granulation tissue, hypodermis, panniculus carnosus muscle, subcutaneous fat pad, fascia and scab (Treuting et al. 2012). We generated an interactive PDF of each scanned H&E image that allows hovering to identify the (x,y) coordinates of each tile (**Fig. 5A**). Using a lasso tool, we generated bounding lines to denote and color code each of the tissue categories (**Fig 5B-D**). Using the multispectral image of the section, we overlaid the color-coded tissue boundaries to verify assignment (**Fig 5E-G**). We performed similar verification of anatomical tissue boundaries using the autofluorescence channel only, as this channel allows visualization of the complete tissue on the slide (**Fig 5H-J**). Next, we applied the color-coded tissue category mask to the tile map and generated tissue category “superclusters” to evaluate distribution of image tiles to each of the 8 categories (**Fig 5K-M**). Using this approach, the number of tiles for scab in WT decrease from POD5 to 10, with concurrent increase in granulation and dermal tissue, as well as in the epidermis (**Fig 5O**). In db/db wounds, scab tiles increase from POD5 to POD10, with notable increase in granulation tissue. Comparison between WT and db/db demonstrates less granulation tissue in db/db at both time points, but also that db/db increase in granulation tissue is far less than that of WT. Analysis of the percentage of tissue across the 4 biological conditions demonstrates similar relationships, aligning with current knowledge of cutaneous wound biology (**Fig 5P**), particularly the severe delays in the db/db model (Peña and Martin 2024; Galiano et al. 2004).

### Assigning super clusters and sub clusters

Building on the density and mean intensity feature representations described above in the Analysis, we next embedded the tile vectors with UMAP (*n_neighbors* = 25, *min_dist* = 0.1) and preformed K-means clustering across a range of *k* values (5-50). Both feature types capture complementary aspects of the signal (i) spatial prevalence versus (ii) per-pixel signal strength while controlling for cross channel scale differences, per-channel z-scoring, ensuring that high intensity channels do not dominate the embeddings, while the exclusion of the autofluorescence channel prevents tile-based separation due to artifacts such as background rather than the true biological signal.

Coverage analysis across *k* values revealed that some tissue categories were absent at certain resolutions or fragmented across multiple clusters, rather than consistently forming distinct groups **(Fig. 6A-B)**. To address this, we merged related groups (hair follicles, interfollicular epidermis, and migrating epithelial tongue into “epithelium”; hypodermis and fat pads into “white adipose tissue, WAT”) which improved overall coverage (Fig. 6C-D). Representative maps illustrate how density-versus intensity-based clustering favored slightly different *k* values (e.g., *k* = 14 for density and *k* = 21 for intensity; Fig. 6E–F). Additional subclustering resolved stromal compartments, yielding the final categories in Fig. 6H–J, which were further characterized by feature bar plots and radar plots (Fig. 6I–L).

**Figure 6.**
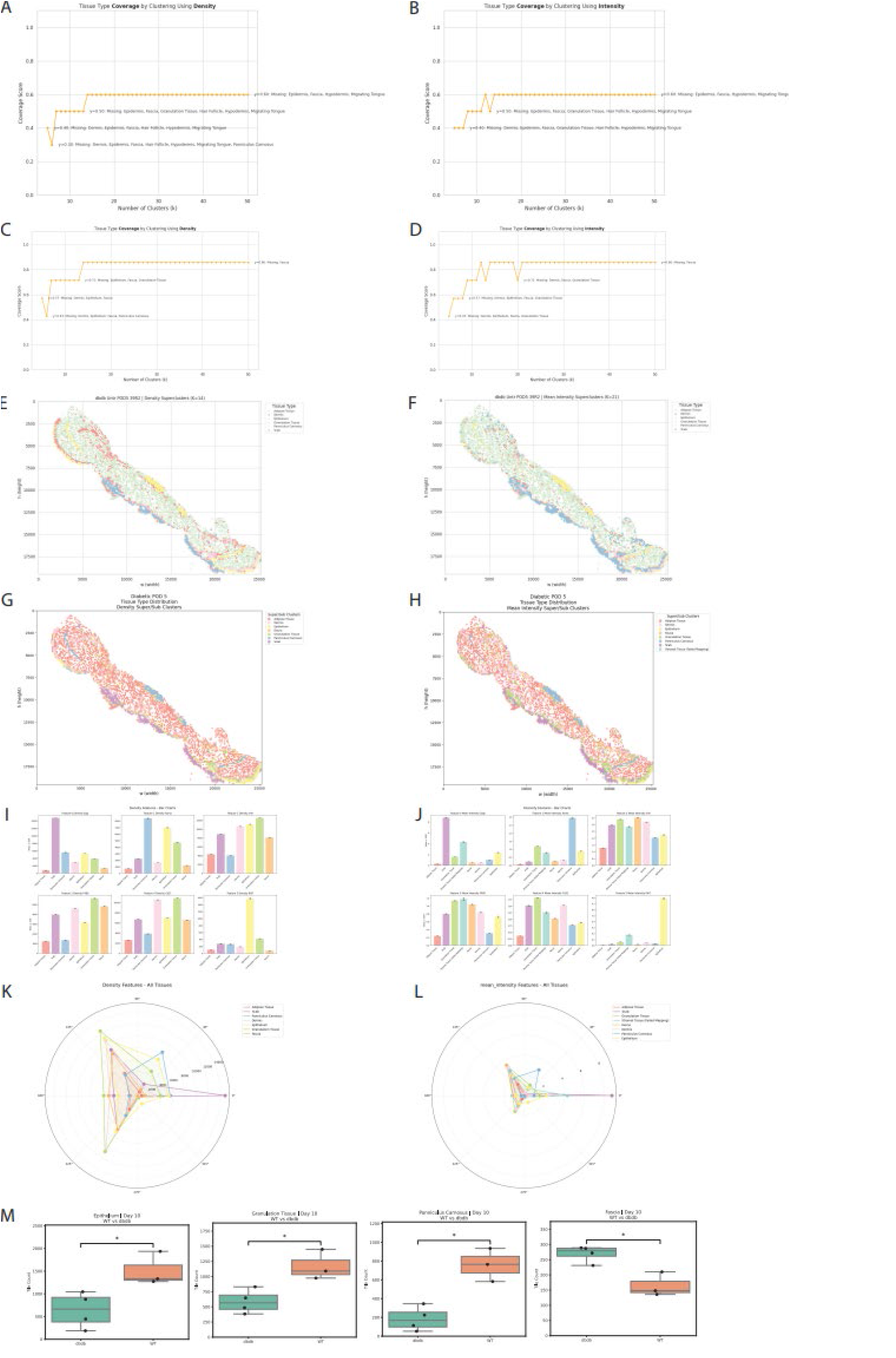
Characterizing the identities of unsupervised clusters and assigning them to cluster groups. (A-B) Coverage score plotted at each K-means clustering level for both density (A) and mean intensity (B). For each of the different Y levels, the tissue type categories which are missing clusters are detailed in the plot. (C-D) Coverage scores recalculated after reducing the number of categories through merger of the above missing tissue types. (E-F) Tile position maps where tiles are colored to depict (E) super clusters calculated using density features at K-means = 14, (F) super clusters calculated using mean intensity features at K means = 21, (G) super clusters calculated using density features at K-means = 14 with subclustering of the stromal tissues. (H-J) Tile position maps where tiles are colored to depict the final categories of clusters that we arrive at using both the super clustering and the subclustering of the stromal tissues. (H) super clusters calculated using mean intensity features at K means = 21 with subclustering of the stromal tissues. (I-L) Plots depicting the final set of clusters and their relationship to the features. (I-J) bar plots for each feature showing its quantification across each of the clusters using (I) density and (J) mean intensity. (K-L) Radar plots showing each of the clusters and how the features define the cluster using (K) density quantifications and (L) mean intensity quantifications. (M) Box plots showing comparing the number of tiles within clusters of wild-type and diabetic samples at POD 10.

Bar plots and radar plots further characterize these categories **(Fig. 6I–L)**. The bar plots reinforce biological validity by showing that cluster identities align with recognizable tissue layers, while also revealing systematic differences in the relative abundance of compartments across experimental groups (e.g., tile number/area differences between WT and db/db wounds at specific timepoints). For each tissue type, significance of genotype differences was assessed using Welch’s two-sample t-tests on per-sample proportions, with thresholds denoted as p < 0.05 (), p < 0.01 (), p < 0.001 (), and n.s. for not significant. In contrast, the radar plots capture the feature-level signatures of each cluster, illustrating how different channels (e.g., CD31, F4/80, αSMA) contribute to defining distinct tissue categories. Together, these visualizations demonstrate not only that clusters reflect anatomically coherent compartments, but also that they are underpinned by biologically interpretable signal characteristics.

## DISCUSSION

Many biological studies rely on accurate segmentation of cell bodies, membranes, and nuclei from microscopy images to quantify tissue architecture and cellular organization (Schindelin et al. 2012; Aeffner et al. 2019). This task becomes particularly challenging when working with large, heterogeneous samples such as whole-tissue sections, where variations in staining quality, tissue morphology, and imaging conditions can introduce significant noise (Gurcan et al. 2009). These challenges are further compounded in pathological contexts, such as diabetic wound healing, where altered vascularization, inflammatory responses, and extracellular matrix remodeling produce complex, atypical signal patterns. These issues are exacerbated by the sampling bias of traditional microscopy.

Whole slides scanning technology addresses this limitation, it eliminates this sample bias by using the power to systematically acquire a full image series across an entire slide. Moreover, the automation of this process significantly reduces the time and labor required for manual microscopy and analysis. Despite these advantages, this relatively recent advent of automated whole-slide scanning presents its own set of challenges which are still being addressed.

Whole-slide scans are substantially larger than those from conventional microscopy, often requiring specialized file formats that differ from the standard TIFF format for processing. These large files pose difficulties for commonly used software that were not originally designed to process whole-slide datasets. Due to the large pixel dimensions and data sizes of whole-slide images, advanced compression techniques and structured metadata are essential for efficient storage, transfer, and analysis. This has led to the adoption of the OME-TIFF format, which builds upon image pyramid structures to manage high-resolution data effectively. However, even with such formats such as OME-TIFF, both file and pixel dimensions still constrain the performance of widely used open-source platforms such as ImageJ/FIJI.

Built on Java, the architecture imposes a strict limitation on maximum array dimensions, specifically that arrays cannot exceed approximately 2^31^ − 1 (≈2.14 billion) elements (elements = width × height × channels × frames) (Rueden et al. 2017; Deroulers et al. 2013). For a single-channel image, this translates to a maximum size limit of 46,000 × 46,000 pixels. When considering a four-channel image (typical for IF images) the maximum size limit is even less (23,000 × 23,000 pixels). Whole slide scans frequently surpass this threshold, commonly exceeding these limits due to extremely large pixel dimensions (often exceeding tens of thousands per axis), multiple imaging channels, and high bit-depth formats (Deroulers et al. 2013). Most 20× objectives produce an effective pixel size around 0.45–0.5 µm/pixel on whole-slide scanners (our data are 0.49 µm/pixel). When considering that the size of a standard microscope slide is roughly ∼ 25mm x 75mm, it can be estimated that a single-channel of a whole slide scan imaged at 20x would have 7.5 billion pixels in total, which is already larger than the 2.14-billion limit on elements of an image. In our study, we have whole slide images obtained with a 20x objective at 32-bit with 7 channels. When we try opening our images using ImageJ/FIJI, the BioFormats plugin attempts to buffer entire image planes, under these conditions, Java’s array limits are breached, often triggering NegativeArraySizeException or ArrayIndexOutOfBoundsException during image opening, even with ample memory allocations (Gatenbee et al. 2023). This indicates that the requested array size exceeds Java’s hard limit rather than signaling file corruption (Deroulers et al. 2013). Consequently, our attempt of increasing ImageJ/FIJI’s memory usage limit 25 GB was insufficient to avoid such crashes. The only workarounds often involve down-sampling, cropping, or bit-depth reduction, which inevitably diminish spatial resolution and compromise quantitative fidelity. Otherwise, researchers are often forced to analyze limited subsets of the data through creating several “high power fields” for their analysis, however this method then reintroduces another form of sampling bias. Thus, the technical barriers to handling whole-slide data are substantial. Yet overcoming these issues only reveals a second, equally important challenge: the analytical frameworks themselves.

Most techniques to analyze whole-slide scans remain rooted in segmentation-based strategies, which were optimized for cultured cells rather than intact tissue architecture. However, these approaches were first created and built using images of in vitro cultured cells and do not recapitulate similar functionality for tissue sections (Zhang et al. 2022; Carpenter et al. 2006). In vitro cultured cells generally have clearly stained and visible nuclei with the entirety of their cell body clearly visible as cultured cells are not situated deeply inside of an organ where they generally more spread out within a tissue matrix. Furthermore, cultured cells usually grow to have a typical, almost standard morphology, again lending to the use of standardized image analysis techniques. Segmentation-based approaches in many cases are significantly more difficult to deploy for images taken from organ specimens. Organs of adult mammals are not translucent in the way that zebra fish embryos are. Structurally, mammalian organs can be composed of multiple tissue types, each with a unique matrix of structural proteins and ECM. These features can significantly complicate technical aspects such as efficacy of antibody based staining and accurate/precise detection of signal from fluorescent proteins.

Organ and tissue structures introduce significant complexity to segmentation-based analysis at the cellular level. Cellular morphologies vary widely across tissue types within an organ, which limits the applicability of cytoplasmic segmentation alone or in combination with nuclear segmentation. In vivo, cells are embedded within non-standardized three-dimensional architectures. As a result, reliable segmentation requires careful tuning of tissue-specific parameters.

Variability in tissue composition further complicates segmentation. Nuclear segmentation is effective only when nuclei are present in the z-frame of the image. That is, when the nuclei lie within the focal plane under analysis. Because mammalian organs are three dimensional, not all cells have nuclei visible in that imaging plane. In some cases, nuclei may be completely absent from the section. Cells without nuclei in the zframe may then be misclassified as background, which leads to quantification errors. In densely packed regions, overlapping nuclei pose an additional problem. Watershed techniques are commonly used to separate clustered nuclei; however, these approaches often cause over-segmentation, where a single nucleus is split into multiple regions, or under-segmentation, where several adjacent nuclei are merged into one.

When nuclear segmentation is not feasible, analysis must rely on cytoplasmic features, but this strategy carries its own risks. Without a visible nucleus, stained cytoplasm may still be mistakenly removed as background. This issue is especially problematic in tissue samples where cellular morphology is not standardized. Specialized cell types exhibit diverse shapes and spatial organization in three dimensions, in contrast with cultured cells, and this disparity increases analytical complexity.

Additional factors further reduce segmentation accuracy. Variation in size, shape, texture and spatial distribution of cellular structures provides important diagnostic information, particularly in cancer detection. Moreover, tissue sectioning thickness and slide mounting methods can alter cytoplasmic and nuclear appearance. These changes may introduce inconsistencies that mimic pathological abnormalities. Noise from stain variation, scanner artifacts and inconsistent contrast adds further difficulty. Another key factor to analysis of this magnitude is specialized cells which can exacerbate these challenges.

Both blood and lymphatic endothelial cells are notoriously known for being one of the most difficult cell types for nuclear segmentation methods. This is because endothelial cells often have thin, elongated structures with nuclei located outside the imaged section. Peripheral nerves are known to send abundant projections into organs, while their cell bodies which house their nuclei will be nowhere near the organs as they are often located around the spinal cord. Lastly immune cells common create issues with cell segmentation of tissue sections. Their circulatory nature makes them abundant in most mammalian organs and they come in a wide range of different types of morphologies. Specifically, macrophages are wildly regarded as being difficult in tissues using microscopy as they display irregular, complex morphologies, and portions of their cell bodies may be visible in a section without the nucleus.

In this work, we circumvent these limitations by leveraging whole-slide multiplex imaging **(Fig. 3)** to perform unsupervised learning directly on spatially registered tile features using a segmentation free approach. Unlike supervised approaches, unsupervised learning has no predefined output set, allowing the algorithm to identify the natural patterns present in the data without prior labels (Komura et al. 2025) By extracting per-tile quantitative measurements from the entire slide, we were able to cluster tiles back onto the tissue map, producing spatially coherent groupings of biologically distinct regions (Fig6. A-D). Unsurprisingly, certain types of tissue, such as scab and adipose tissue, were readily distinguishable. Tiles containing scab typically exhibited high cellular density and complex surface morphology, whereas adipose tiles contained sparse nuclei and characteristic large, low-intensity vacuoles. While such pronounced phenotypic contrasts made the separation in feature space biologically unsurprising, it is noteworthy that these patterns emerged so clearly from a fully unsupervised analysis of whole-slide tile features. As shown in **Fig. 4**, even before clustering, tile-level intensity distributions and coverage rankings already hinted at systematic differences between tissue types and biological conditions. In line with these patterns, **Fig. 6**, stark contrasts like scab versus adipose naturally separated in feature space, confirming that the extracted features captured meaningful biological variation. Beyond these obvious distinctions, the same approach also delineated more subtle compartments, including epidermis and stromal layers. In the super clustering UMAP space (**Fig. 6**), these tissues appeared as neighboring clusters rather than being widely separated, reflecting their closer phenotypic similarity while nonetheless remaining consistently distinguishable.

Importantly, because this analysis is applied across all tiles within each slide, it captures quantitative variation at a scale that enables direct comparison between biological conditions. Each tile spans ∼63.5 × 63.5 µm (F**ig. 4A– E**), which is large enough to encompass multiple cell types and microstructural features of interest. In densely cellular regions such as scab or epidermis, a tile typically contains dozens of cells, including clusters of macrophages, while in sparser regions such as adipose tissue it may contain only a few nuclei surrounded by large lipid vacuoles **(Fig. 4**). At this scale, vascular structures, particularly small vessels and capillary segments marked by CD31, can also be fully contained within a single tile, enabling co-localization analysis of immune and endothelial markers. This per-tile granularity allows aggregate distributions to reflect both cellular composition and spatial organization. Using the same unsupervised feature space, we were able to separate diabetic wound models from wild-type controls based on these aggregate tile-level distributions or area **(Fig. 6)**. This demonstrates that, beyond resolving obvious tissue compartments, our approach detects subtle yet systematic differences in tissue organization and composition associated with disease state.

In this study, we deliberately combined manual annotation with unsupervised clustering to both validate and interpret our findings. Prior annotation of wound sections (**Fig. 5**) established a biological reference frame, ensuring that major compartments such as scab, adipose, epidermis, and stroma were clearly defined in their spatial context. The subsequent unsupervised clustering reproduced these tissue categories directly from tile-level features, with cluster identities confirmed through annotation of representative samples. Importantly, while these manual labels served as interpretive anchors, the clustering itself was not driven by them. The ability of unsupervised analysis to recapitulate known tissue compartments, while also resolving their spatial relationships within wound sections, highlights the method’s capacity to uncover biologically meaningful structure without prior labels.

In sum, our work introduces the framework for unbiased analysis of WSIs, revealing how segmentation free, tile-based clustering can resolve the cellular and structural hallmarks of healing. By employing ML rather than deep neuronal networks, our approach (i) maintains interpretability and reduces computational demands, (ii) and remains robust in settings with limited training data, an advantage for biological imaging, where annotated dataset is over scare or labor intensive to generate (Sharma et al. 2021). This offers a broadly applicable framework for studying regenerative pathology in mammalian systems, while also providing insight into disease associate models.

